# Redefining the *Clostridioides difficile* σ^B^ regulon: σ^B^ activates genes involved in detoxifying radicals that can result from the exposure to antimicrobials and hydrogen peroxide

**DOI:** 10.1101/2020.07.16.207829

**Authors:** Ilse M. Boekhoud, Annika-Marisa Michel, Jeroen Corver, Dieter Jahn, Wiep Klaas Smits

## Abstract

In many gram-positive bacteria the general stress response is regulated at the transcriptional level by the alternative sigma factor sigma B (σ^B^). In *C. difficile* σ^B^ has been implicated in protection against stressors such as reactive oxygen species and antimicrobial compounds. Here, we used an anti-σ^B^ antibody to demonstrate time-limited overproduction of σ^B^ in *C. difficile* despite its toxicity at higher cellular concentrations. This toxicity eventually led to the loss of the plasmid used for anhydrotetracycline-induced σ^B^ gene expression. Inducible σ^B^ overproduction uncouples σ^B^ expression from its native regulatory network and allowed for the refinement of the previously proposed σ^B^ regulon. At least 32% the regulon was found to consist of genes involved in the response to reactive radicals. Direct gene activation by *C. difficile* σ^B^ was demonstrated through *in vitro* run-off transcription of specific target genes (*cd0350, cd3614, cd3605, cd2963*). Finally, we demonstrated that different antimicrobials and hydrogen peroxide induce these genes in a manner dependent on this sigma factor, using a plate-based luciferase reporter assay. Together, our work suggests that lethal exposure to antimicrobials may result in the formation of toxic radicals that lead to σ^B^-dependent gene activation.

**Importance:** Sigma B is the alternative sigma factor governing stress response in many gram-positive bacteria. In *C. difficile*, a *sigB* mutant shows pleiotropic transcriptional effects. Here, we determine genes that are likely direct targets of σ^B^ by evaluating the transcriptional effects of σ^B^ overproduction, provide biochemical evidence of direct transcriptional activation by σ^B^, and show that σ^B^-dependent genes can be activated by antimicrobials. Together our data suggest that σ^B^ is a key player in dealing with toxic radicals.

## Introduction

Disruption of the normal gastrointestinal flora as a result of antimicrobial treatment can lead to a *Clostridioides* (*Clostridium*) *difficile* infection (CDI) (1). *Clostridoides difficile* is a gram-positive, spore-forming obligate anaerobe and the primary cause for nosocomial infectious diarrhea (2). Its highly resistant endospores are usually transmitted via the oral-fecal route and germinate into vegetative cells in the colon upon contact with primary bile acids and other inducing factors (3). In the gut vegetative *C. difficile* cells are facing many environmental stressors, including variations in oxygen tension, pH, osmolarity, nutrient availability, and the inflammatory responses of the immune system (4). The bacteria are also faced with antimicrobial compounds produced by the host, the resident microbiota, or given externally during medical therapy (5). The physiological response of *C. difficile* to these insults and the inflammatory responses triggered by CDI can result in the production of Reactive Oxygen Species (ROS), Reactive Nitrogen Species (RNS) and nitric oxide (NO) (2, 6). Bacteria need to adapt to changing environmental conditions, including stresses, by adapting their physiology in a timely manner. This is achieved by fast transcriptional reprogramming, followed by shortly delayed changes at the translational level (7). The alternative sigma factor sigma B (σ^B^, encoded by the *sigB* gene) - which regulates the general stress responses in a variety of gram-positive organisms - is central to the maintenance of the cellular homeostasis during stress adaptation (8, 9).

Sigma factor B activity in Firmicutes is regulated at the protein level by a partner-switching mechanism in which the anti-sigma factor RsbW binds and inhibits σ^B^ association with the RNA polymerase under non-stressed conditions. When a σ^B^-activating stress is sensed the anti-anti-sigma factor RsbV is dephosphorylated by a phosphatase. Subsequently, RsbV is able to bind and sequester the anti-sigma factor RsbW, allowing for the association of free σ^B^ with the RNA polymerase core enzyme (8, 10). The factors ultimately involved in sensing and the corresponding dephosphorylation of RsbV are stressor- and species-specific. For instance, *Bacillus subtilis* and *Listeria monocytogenes* respond to environmental stresses through a stressosome composed of RsbR/S/T/X, encoded by genes upstream of *rsbV, rsbW* and *sigB* in the same operon (8). Ultimately, this leads to activation of the RsbU phosphatase and dephosphorylation of RsbV (8). The *C. difficile sigB* operon does not contain genes encoding RsbR/S/T/X homologs. Two open reading frames upstream of *rsbV* (*cd0007*-*cd0008*) appear to have no role in σ^B^ activation (11). Recently a phosphatase, RsbZ, responsible for RsbV dephosphorylation has been characterized in *C. difficile*, but the gene encoding this protein is not a part of the *sigB* operon (11). Tight regulation of σ^B^ by the partner-switching mechanism is required as the energy burden associated with σ^B^ activity was found to be disadvantageous in several different organisms (12, 13). Nevertheless, σ^B^ is essential for in-host survival for several pathogenic bacterial species. For example, in *L. monocytogenes* σ^B^ is involved in counteracting the effects of the acidic pH encountered in the stomach and upon invasion of intestinal epithelial cells in the lysosome (14, 15). Antimicrobial resistance in certain pathogenic bacteria is σ^B^-dependent. In *Staphylococcus aureus* σ^B^ overproduction leads to thickening of the cell wall and increased resistance to beta-lactam antimicrobials (16). The *sigB* homologue *sigF* of *Mycobacterium tuberculosis* is induced by small amounts of rifamycin (17). Analogously, *B. subtilis* σ^B^ is involved in resolving a rifampin-induced growth arrest (18). There is also evidence for the involvement of σ^B^ in *C. difficile* in the response to antimicrobial substances. Mutants of *sigB* show increased susceptibility to rifampicin and mitomycin C, and are also more sensitive to hydrogen peroxide, nitroprusside and di-ethylamine NONOate. However, the underlying molecular mechanism remains unknown (19). Finally, indirect activation of σ^B^-dependent genes as the result of a gene-dosage shift has been demonstrated for *C. difficile* exposed to DNA-polymerase inhibitors such as the Phase I drug ibezapolstat/ACX-362E (20). In this study, we demonstrate that despite its detrimental cellular effects σ^B^ overexpression is detectable and tolerated for short periods of time. This allowed for the experimental identification of a set of genes that is most likely directly regulated by σ^B^. The obtained results show that genes involved in the oxidative and nitrosative stress response form the core of the regulon. Additionally, we show that various antimicrobials and hydrogen peroxide induce the expression of σ^B^-regulated genes in a σ^B^-dependent manner, suggesting a link between the lethal exposure to antimicrobials and oxidative and nitrosative stresses in *C. difficile*.

## Results

### C. difficile σ^B^ is measurably overproduced upon induction of the sigB gene

Previous investigations of σ^B^ in *C. difficile* have used a *sigB* mutant and characterized its gene expression in the stationary growth phase in comparison with a wild type strain. Though informative, this method is likely to result in indirect effects of σ^B^ due to stationary phase heterogeneity, prolonged incubation and possible positive or negative feedback in the σ^B^ regulatory circuit. To circumvent these issues and identify genes likely to be regulated by σ^B^ directly, we set out to uncouple *sigB* expression from its native regulatory circuit by expressing it from an inducible promoter.

Firstly, in order to confirm overproduction of σ^B^, we measured cellular σ^B^ levels using immunoblotting. For this purpose, we heterologously overproduced and purified σ^B^ containing a C-terminal His-tag (**Figure 1A**) and used this protein to raise a polyclonal antiserum. Corresponding polyclonal antibodies were affinity-purified to prevent unspecific immune reactions.

**Figure 1.**
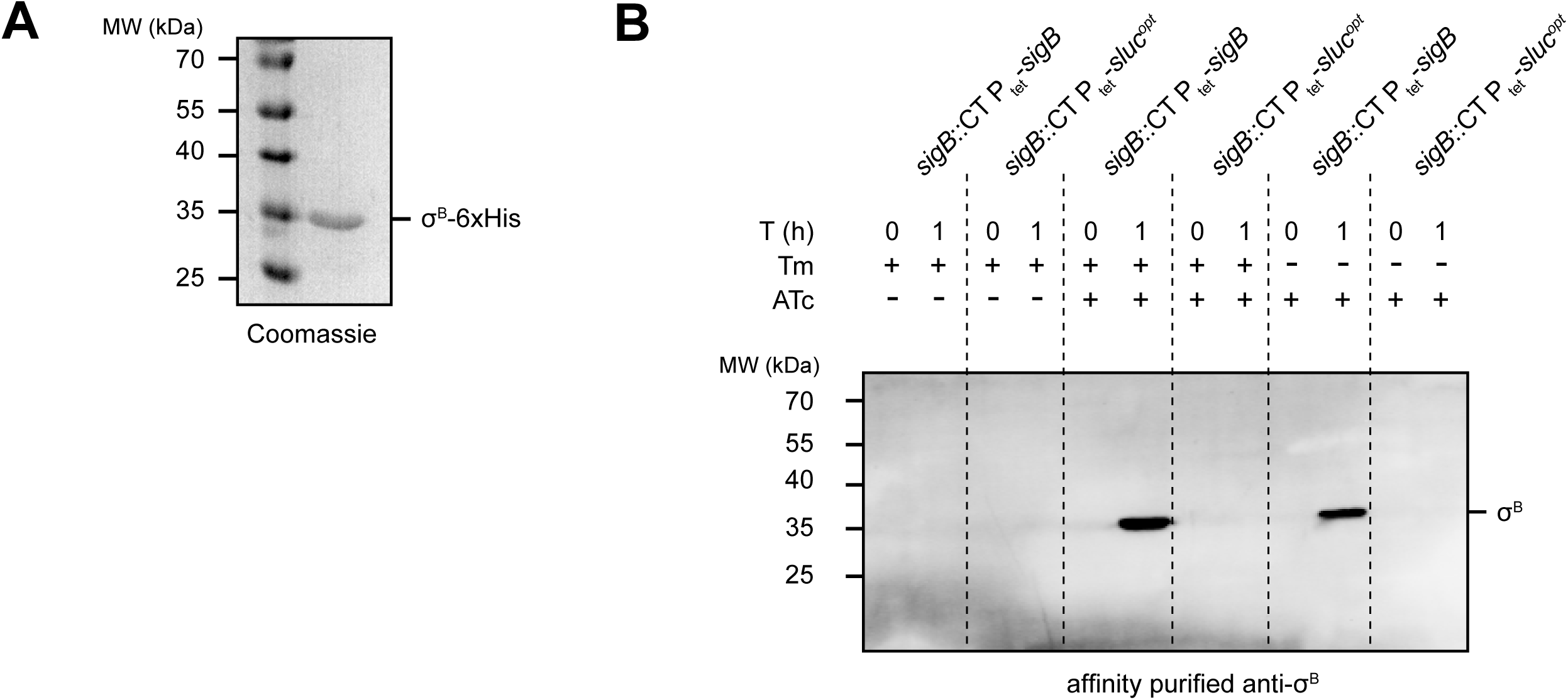
Recombinant σ^B^ was used to generate a *C. difficile* specific antibody for intracellular detection. **A**) Coomassie Blue-stained 12.5% SDS-PAGE gel of purified recombinant σ^B^-6xHis. **B**) Western Blot using affinity purified σ^B^ antibody (1:500) on strains IB58 (*sigB*::CT P_tet_--*sigB*) and IB61 (*sigB*::CT P_tet_-*sluc*^*opt*^). Cells were grown in lincomycin (20 µg/mL) and in the presence or absence of thiamphenicol (20 µg/mL) until OD_600nm_≈0.3, after which the indicated samples were induced with 100 ng/mL ATc. Samples were collected directly after the addition of ATc (or at the time ATc would have been added in the uninduced controls) at T=0h and after 1 hour of induction (T=1).

Next, we set out to validate the overproduction of σ^B^ in transconjugant *C. difficile* cells harboring plasmids containing *sigB* under the control of the anhydrotetracyclin (ATc)-dependent promoter P_tet_ (21). For this purpose, σ^B^ was produced in a *sigB* mutant background (strain IB58; *sigB*::CT P_tet_-*sigB*). As a control, we introduced a non-related expression construct in the same strain (IB61; *sigB*::CT P_tet_-*sluc*^*opt*^).

We expected a signal at approximately 30 kDa in Western blot experiments for cells grown in the presence of inducer ATc for strain IB58, but not for the uninduced cultures of IB58 or the control strain IB61. Additionally, by growing in the presence or absence of thiamphenicol, we investigated whether overproduction of σ^B^ required selection for the P_tet_-*sigB* expression plasmid.

When strains were grown in BHIY medium supplemented with 20 µg/mL lincomycin and induced for one hour with or without 100 ng/mL ATc in the presence or absence of 20 µg/mL thiamphenicol, we did not detected any signal at the molecular weight expected for σ^B^ in the ATc-induced control samples (*sigB*::CT P_tet_-*sluc*^*opt*^) or in any of the uninduced samples (**Figure 1B**). In contrast, after one hour of induction a clear band of the expected molecular weight of σ^B^ (∼30 kDa) was observed only in the IB58 (*sigB*::CT P_tet_-*sigB*) samples (**Figure 1B**). Plasmid selection by inclusion of thiamphenicol in the growth medium did not influence σ^B^ overproduction in this time frame.

We conclude that the affinity purified rabbit-α-σ^B^ antibody is specific for σ^B^ and can be used for its detection in lysates of *C. difficile*. Furthermore, it is possible to uncouple *sigB* expression from its tight regulatory network by ATc-inducible overexpression for one hour *in trans*.

### Prolonged overexpression of σ^B^ is lethal and leads to loss of plasmids harboring P_tet_-sigB

Above we showed that it is possible to overproduce σ^B^ in *C. difficile* and that it is tolerated by the bacterium for one hour. This observation is somewhat at odds with the previously reported toxic nature of overproduced σ^B^ (8, 11). To reconcile these two observations, the effect of long-term overexpression of *sigB* and the stability of the plasmids used for σ^B^ overproduction under such conditions were investigated. First, overnight cultures of 630Δ*erm* (wildtype), AP34 (P_tet_-*sluc*^*opt*^) and JC096 (P_tet_-*sigB*) were adjusted for their OD_600nm_ values and ten-fold serially diluted. Subsequently, 2 µl spots per dilution were made on selective (20 µg/mL thiamphenicol) and non-selective BHIY agar plates, part of which contained 200 ng/mL ATc to induce P_tet_-dependent gene expression. All plates were then incubated anaerobically for 24 hours. On plates without thiamphenicol, regardless of the presence of the inducer ATc, comparable growth was observed for all three strains (**Figure 2A**). As expected, when selecting for the plasmid using thiamphenicol no growth was observed for the susceptible 630Δ*erm* strain (which lacks the *catP* gene contained on the expression vector). In the absence of the inducer, no difference in growth was observed for the vector control strain (AP34; P_tet_-*sluc*^*opt*^) compared to the strain carrying the P_tet_-*sigB* plasmid (JC096). However, upon induction of *sigB* expression on selective plates a 3-4 log growth defect was observed for the strain carrying P_tet_-*sigB* compared to the vector control strain. We conclude that prolonged induction of *sigB* expression is toxic when cells are cultured in the presence of thiamphenicol. Our results thus corroborate the finding that σ^B^ overproduction is toxic to *C. difficile* cells in liquid culture (11).

**Figure 2.**
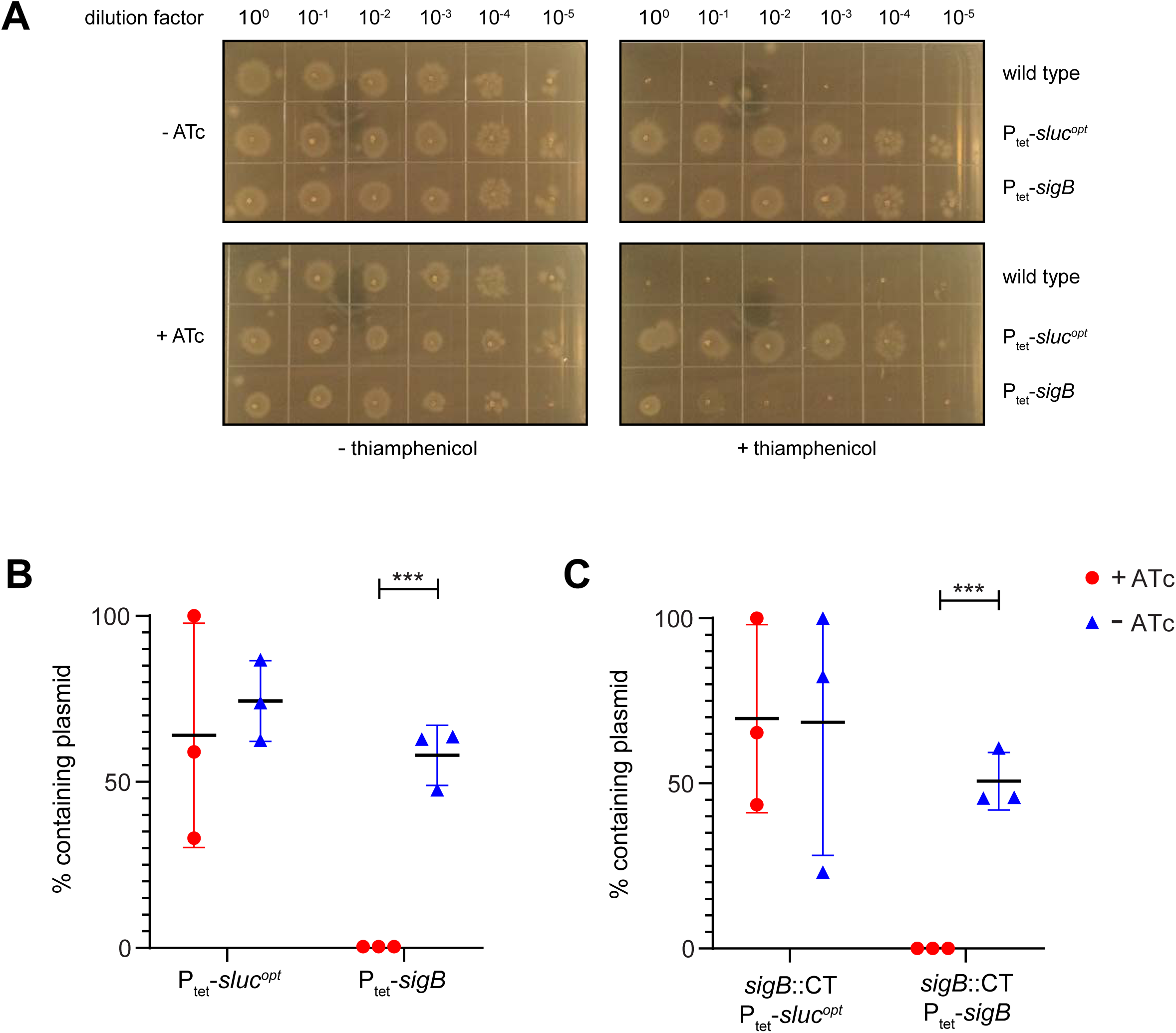
Overexpression of σ^B^ is toxic for *C. difficile* and leads to plasmid loss. **A**) Tenfold serial dilutions on BHIY agar plates of 630Δ*erm* (wildtype), AP34 (P_tet_-*sluc*^*opt*^) and JC096 (P_tet_-*sigB*). Similar results were obtained for strains IB58 and IB61 (data not shown). **B**) Percentage of cells retaining the plasmid in AP34 (P_tet_-*sluc*^*opt*^) and JC096 (P_tet_-*sigB*). **C**) Percentage of cells retaining the plasmid in strains IB61 (*sigB*::CT P_tet_-*sluc*^*opt*^) and IB58 (*sigB*::CT P_tet_-*sigB*). Percentages were calculated based on the ratio of CFU/mL of the paired selective (with thiamphenicol)- and non-selective (without thiamphenicol) plates. *** p≤0.001 as determined by an unpaired student’s t-test, n=3.

The lethality associated with σ^B^ overproduction was not seen when cells were grown without thiamphenicol in our experiment (**Figure 2A**). We considered two possible explanations for this observation. As thiamphenicol is used for ensuring plasmid maintenance, its absence might result in plasmid loss when a toxic protein such as σ^B^ is overproduced. The remaining cells, that no longer express σ^B^ would consequently be susceptible to thiamphenicol (due to the loss of *catP*). Alternatively, the combination of σ^B^ and thiamphenicol might be toxic to the bacteria. To test whether plasmid loss was the cause of the observed lethality of bacteria overproducing σ^B^ in the presence of thiamphenicol, cells from the plates without thiamphenicol (with and without ATc) were resuspended in BHIY medium, adjusted for their OD_600nm_ and ten-fold serially diluted. Ten-µl spots of these dilutions were plated on plasmid-selective (thiamphenicol) and non-selective (no thiamphenicol) plates. Based on the ratio of CFU/mL of the selective and non-selective plates, the percentage of cells which lost their plasmid was calculated. If σ^B^ overproduction led to the loss of the plasmid under conditions that do not select for its maintenance (no thiamphenicol), we expected significantly reduced growth on plates containing thiamphenicol. Although some plasmid loss was observed under uninduced conditions, as well as for the negative control strain AP34 (P_tet_-*sluc*^*opt*^)(**Figure 2B**), all cells originally containing the P_tet_-*sigB* plasmid (strain JC096) lost this plasmid upon induction of σ^B^ overproduction with ATc. Similar results were obtained for *sigB* mutant strains IB58 and IB61 (**Figure 2C**), indicating the observed effects were solely due to *in trans* σ^B^ overproduction, and did not result from an interference of the native *sigB* regulatory network. Together, these results are consistent with a model in which the vector with the low-copy number pCD6 replicon are rapidly eradicated upon expression of a gene (here *sigB*) that causes lethal defects (22, 23).

### σ^B^ primarily activates genes relating to oxidative/nitrosative stress responses

Above, we have shown that long-term overproduction σ^B^ is detrimental and that this leads to loss of the expression plasmid in the absence of thiamphenicol (**Figure 2**), but that σ^B^ overproduction nevertheless could clearly be demonstrated when induction is limited to 1 hour (**Figure 1C**). Therefore we used the time-limited induction to refine the previously proposed regulon (19) in both the presence and absence of thiamphenicol to strike a balance between potential secondary effects due to toxicity associated with σ^B^ overproduction (with thiamphenicol), and loss of the expression plasmid from a subpopulation of cells (without thiamphenicol)(**Table 1**). We compared transcriptome data from strain IB58 (*sigB::*CT P_tet_-*sigB*) to that of strain IB61 (*sigB::*CT P_tet_-*sluc*^*opt*^). IB61 harbors a vector for the inducible expression of a luciferase gene which does not lead to any toxicity and growth phenotype (24)

**Table 1.** Setup of the DNA array and number of differentially expressed (DE) genes, including number of positively (POS) and negatively (NEG) regulated genes. Numbers inbetween brackets correspond to the number of differentially expressed genes after the 5 genes from hybridisations 1 and 3 are excluded.

|  | Control | Target | Conditions | DE | POS | NEG |
| --- | --- | --- | --- | --- | --- | --- |
| **1** | *sigB*::CT P<sub>tet</sub><sup>-</sup><br>*sluc*<sup>opt</sup> (IB61) | *sigB*::CT<br>P<sub>tet</sub>-*sigB*<br>(IB58) | Lincomycin [20µg/mL],<br>Thiamphenicol [20µg/mL],<br>no ATc | 5 | 4 | 1 |
| **2** | *sigB*::CT P<sub>tet</sub><sup>-</sup><br>*sluc*<sup>opt</sup> (IB61) | *sigB*::CT<br>P<sub>tet</sub>-*sigB*<br>(IB58) | Lincomycin [20µg/mL],<br>Thiamphenicol [20µg/mL],<br>ATc [100 ng/mL] | 183 (178) | 167<br>(163) | 16 (15) |
| **3** | *sigB*::CT P<sub>tet</sub><sup>-</sup><br>*sluc*<sup>opt</sup> (IB61) | *sigB*::CT<br>P<sub>tet</sub>-*sigB*<br>(IB58) | Lincomycin [20µg/mL], no<br>Thiamphenicol, no ATc | 5 | 4 | 1 |
| **4** | *sigB*::CT P<sub>tet</sub><sup>-</sup><br>*sluc*<sup>opt</sup> (IB61) | *sigB*::CT<br>P<sub>tet</sub>-*sigB*<br>(IB58) | Lincomycin [20µg/mL], no<br>Thiamphenicol, ATc [100<br>ng/mL] | 150 (145) | 136<br>(132) | 14 (13) |

We expected no or a limited number of genes to be differentially expressed (log2 foldchange (log2FC) of ≤ -1.5 or ≥1.5, and adjusted p-value of <0.05) under non-inducing conditions. Indeed, we found only five differentially expressed genes in the P_tet_-*sigB* strain (IB58) compared to the P_tet_-*sluc*^*opt*^ control (IB61) strain (hybridizations 1 and 3) (**Dataset S1**). These genes were similarly positively (CD0583, CD0584, both GGDEF domain containing proteins (25); CD2214, CD2215, both potential transcriptional regulators (26)) and negatively (CD1616, an EAL domain protein (25)) regulated in all hybridizations, including those where *sigB* expression was not induced. These results suggest that the basis for the observed differential expression of these genes was vector specific, but not dependent on σ^B^ induction. These genes were therefore not investigated further and are excluded from the numbers discussed below.

Upon induction of *sigB* expression 145 genes were differentially expressed when strains were cultured without thiamphenicol (hybridization 4), and 178 genes were differentially expressed when thiamphenicol was present during cultivation (hybridization 2)(**Figure 3 and Dataset S1**). The majority showed an increase in expression upon induction of *sigB* expression (132 in the samples without thiamphenicol, and 163 in the samples with thiamphenicol), while a minority revealed a decreased expression (13 in the samples without thiamphenicol, and 16 in the samples with thiamphenicol). Of note, we observed only a minor difference in the number of differentially expressed genes between the cells grown in the absence and presence of thiamphenicol (33 genes).

**Figure 3.**
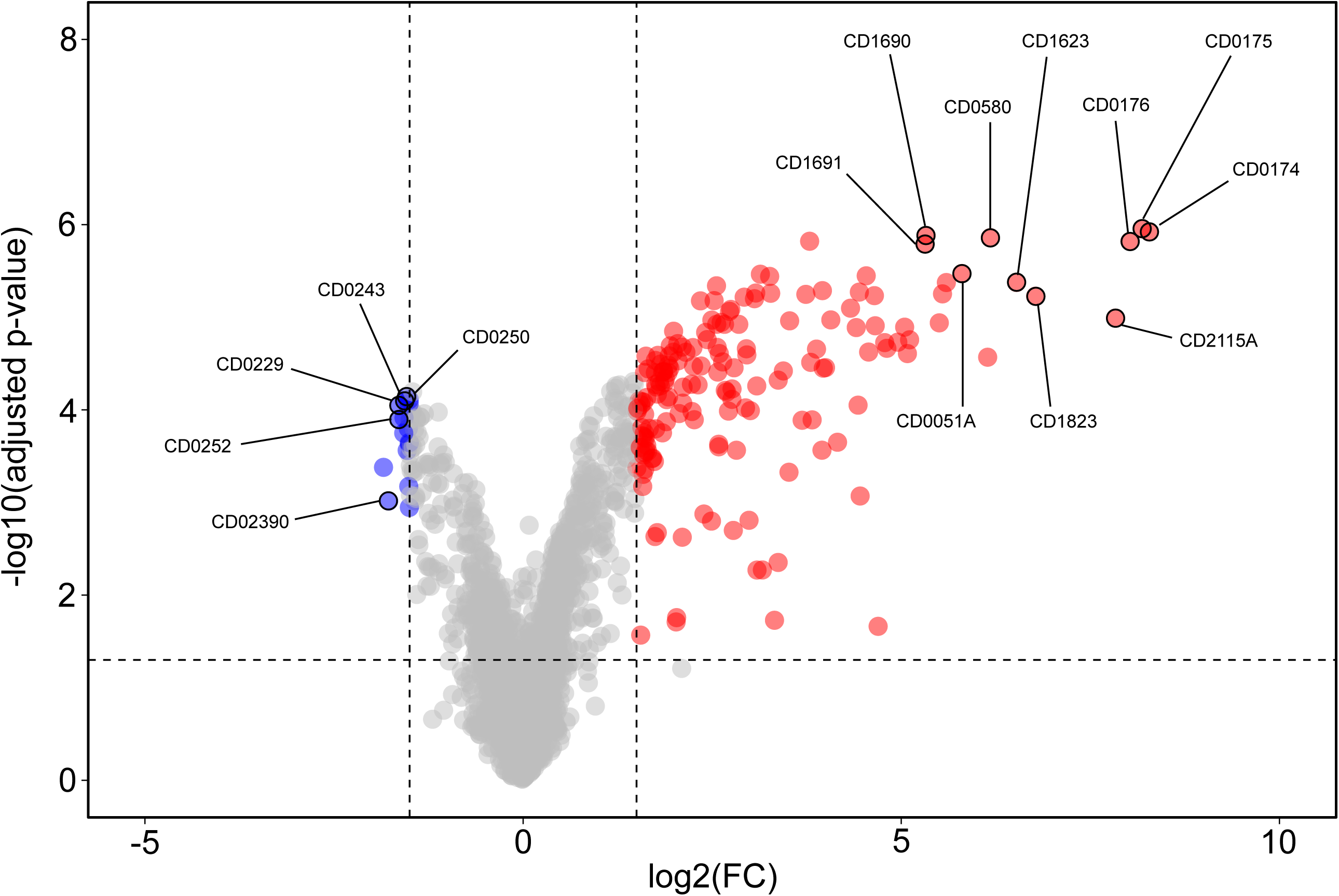
Volcano plot of the transcriptome analysis of the σ^B^ regulon. Graphical representation of differential gene regulation upon overproduction of σ^B^. Dashed lines indicated the significance threshold: |log2FC|>1.5 and adjusted p-value <0.05. Genes significantly up-regulated by σ^B^ are indicated in red, and genes downregulated are indicated in blue. The top-10 of up-regulated genes and 5 selected down-regulated genes are annotated in the figure. An interactive version of the graph is available for exploration via the URL provided in the **Text S1**.

Together, these results demonstrate a high level of consistency in the σ^B^ regulon, despite potential plasmid-loss (when grown in the absence of thiamphenicol) or toxic effects (when grown in the presence thiamphenicol). Our results also show that σ^B^ primarily activates gene expression.

We focused our further analyses on the data obtained from hybridization 2 (with ATc and thiamphenicol), as this condition provided the broadest dataset (178 differentially expressed genes) for the re-definition of the σ^B^ regulon under our experimental conditions (**Dataset S1**).

Of the 163 genes up-regulated by σ^B^ the vast majority appeared to be associated with an response to oxidative stress, since they encode various oxidoreductases, peroxidases, and thioredoxin reductases (**Table 2**). Notably, approximately 51% of the 98 genes previously found to be up-regulated under aerobic stress (7) were also positively regulated by σ^B^ (**Table 2**). Five additional genes associated with aerobic/nitrosative stress (CD0174/*cooS*, CD1279/*iscS2*, CD1280, CD1594/*cysK*, CD1823 and CD2166/*msrAB*) were also found to be induced by σ^B^, in agreement with previous findings (19).

**Table 2.** Selected genes differentially expressed upon overexpression of σ^B^. Genes positively regulated by σ^B^ aerobic stress and other genes involved in oxidative/nitrosative stress (7, 19) are highlighted here. CD numbers corresponding to the published annotation of strain CD630 (52) can be derived by removing “630DERM_” and removing the last digit (in case of a 0) or replacing it with an A (in the case of a 1).

| Locus tag | Gene name | log2FC | adjusted p-value | Predicted function |
| --- | --- | --- | --- | --- |
| **Genes upregulated by aerobic stress and positively regulated by $\sigma^B$** |  |  |  |  |
| CD630DERM_00530 | *mrnC* | 1.9 | 3.29E-05 | Ribonuclease III domain |
| CD630DERM_01750 |  | 8.2 | 1.11E-06 | Oxidoreductase, Fe-S subunit |
| CD630DERM_01760 |  | 8.0 | 1.52E-06 | Oxidoreductase, NAD/FAD binding subunit |
| CD630DERM_01920 | *cls* | 2.6 | 2.35E-04 | Cardiolipin synthetase 1 |
| CD630DERM_03500 |  | 1.9 | 4.63E-05 | Putative hydrolase, HAD superfamily |
| CD630DERM_03510 |  | 1.8 | 5.66E-05 | Conserved hypothetical protein |
| CD630DERM_05600 | *nfo* | 4.0 | 3.5E-05 | Endonuclease IV |
| CD630DERM_05610 |  | 2.0 | 2.39E-05 | Putative aldo/keto reductase; putative ferredoxin |
| CD630DERM_05650 | *nth* | 3.7 | 5.66E-06 | Endonuclease III |
| CD630DERM_05660 |  | 2.5 | 6.62E-06 | Putative tRNA/rRNA methyltransferase |
| CD630DERM_05800 | *gapN* | 6.2 | 1.38E-06 | Glyceraldehyde-3-phosphate dehydrogenase (NADP(+)) (GADPH) |
| CD630DERM_11250 |  | 5.5 | 1.15E-05 | Nitroreductase-family protein |
| CD630DERM_11570 | *norV* | 4.0 | 5.16E-06 | Anaerobic nitric oxide reductase flavorubredoxin (FIRd) (FlavoRb) |
| CD630DERM_13410 | *exoA* | 1.8 | 6.76E-05 | Exodeoxyribonuclease |
| CD630DERM_14630 |  | 1.8 | 1.77E-04 | Conserved hypothetical protein |
| CD630DERM_15240 |  | 4.8 | 1.88E-05 | Putative rubrerythrin |
| CD630DERM_15260 | *pyrC* | 1.6 | 1.98E-04 | dihydroorotase |
| CD630DERM_15760 |  | 1.6 | 2.71E-02 | putative arylesterase |
| CD630DERM_16220 |  | 2.7 | 1.03E-04 | Putative lipoprotein |
| CD630DERM_16230 |  | 6.5 | 4.21E-06 | NADH-oxygen oxidoreductase |
| CD630DERM_16240 | *vanR* | 1.6 | 8.84E-05 | two-component response regulator |
| CD630DERM_16900 | *trxA* | 5.3 | 1.31E-06 | Thioredoxin reductase |
| CD630DERM_16910 | *trxB* | 5.3 | 1.62E-06 | Putative thioredoxin-disulfide reductase |
| CD630DERM_17780 |  | 2.6 | 1.19E-05 | Conserved hypothetical protein |
| CD630DERM_17790 |  | 2.4 | 1.46E-05 | Conserved hypothetical protein |
| CD630DERM_18180 | *ispH* | 1.8 | 5.35E-05 | 4-hydroxy-3-methylbut-2-enyl diphosphate reductase |
| CD630DERM_18220 | *bcp* | 6.1 | 2.70E-05 | Putative thiol peroxidase |
| CD630DERM_18970 |  | 1.8 | 3.20E-05 | Conserved hypothetical protein |
| CD630DERM_19430 |  | 3.3 | 3.62E-06 | Conserved hypothetical protein |
| CD630DERM_20460 |  | 5.5 | 5.60E-06 | Conserved hypothetical protein |
| CD630DERM_21170 | *trxB2* | 5.0 | 1.86E-05 | Thioredoxin reductase |
| CD630DERM_21650 |  | 2.2 | 1.04E-04 | Transcriptional regulator, HTH-type |
| CD630DERM_24760 |  | 4.6 | 5.84E-06 | Conserved hypothetical protein |
| CD630DERM_27960 | *cwp10* | 5.1 | 2.47E-05 | Cell surface protein |
| CD630DERM_27970 |  | 5.1 | 1.75E-05 | Putative calcium-binding adhesion protein |
| CD630DERM_29930 |  | 3.8 | 3.07E-05 | Conserved hypothetical protein |
| CD630DERM_30380 |  | 3.9 | 2.21E-05 | Conserved hypothetical protein |
| CD630DERM_30390 |  | 4.1 | 1.07E-05 | Putative ATPase |
| CD630DERM_30400 |  | 2.1 | 8.46E-05 | Conserved hypothetical protein |
| CD630DERM_30420 |  | 2.6 | 2.47E-05 | Putative membrane protein |
| CD630DERM_33070 |  | 3.8 | 1.51E-06 | Putative phosphoesterase |
| CD630DERM_33100 |  | 3.1 | 6.31E-06 | Putative D-isomer specific 2-hydroxyacid dehydrogenase |
| CD630DERM_33110 |  | 2.9 | 1.19E-05 | Conserved hypothetical protein |
| CD630DERM_34080 |  | 2.3 | 3.38E-05 | Putative DNA mismatch repair ATPase MutS |
| CD630DERM_34090 | *scoC/hprK* | 2.4 | 1.74E-05 | PTS system, HPr kinase/phosphorylase |
| CD630DERM_34100 | *uvrC* | 2.7 | 1.19E-05 | Excinuclease subunit C |
| CD630DERM_34730 | *atpC2* | 1.6 | 2.90E-04 | ATP synthase C chain |
| CD630DERM_34760 | *atpZ* | 1.6 | 2.42E-04 | putative ATP synthase protein |
| CD630DERM_36100 |  | 4.6 | 2.38E-05 | Conserved hypothetical protein |
| CD630DERM_36140 |  | 2.5 | 1.07E-05 | Conserved hypothetical protein, DUF1130 family |
| **Other genes involved in oxidative/nitrosative stress positively regulated by $\sigma^B$** |  |  |  |  |
| CD630DERM_01740 | *cooS* | 8.3 | 1.20E-06 | Carbon monoxide dehydrogenase |
| CD630DERM_12790 | *iscS2* | 2.8 | 2.00E-03 | Cysteine desulfurase |
| CD630DERM_12800 |  | 3.0 | 1.56E-03 | Putative NifU-like protein |
| CD630DERM_14740 |  | 5.0 | 1.28E-05 | Putative rubrerythrin (Rr) |
| CD630DERM_15940 | *cysK* | 3.4 | 4.44E-03 | O-acetyl-serine thiol-lyase A (O-acetyl-sulfhydrylase)(OAS-TL) |
| CD630DERM_18230 |  | 6.8 | 5.93E-06 | Conserved hypothetical protein, UPF0246 family |
| CD630DERM_21660 | *msrAB* | 4.7 | 1.24E-05 | Peptide methionine sulfoxide reductase MsrA/MsrB |

Our findings are recapitulated in a volcano plot (27), that clearly shows that genes with lower expression upon *sigB* induction (in blue) cluster close to the significance threshold whereas those with increased expression (in red) show a larger fold change (**Figure 3**). We calculated the Manhattan distance for each data point (**Dataset S2**), and discuss the proteins encoded by the top-10 of the differentially expressed genes below.

CD0051A is a small hypothetical protein of unknown function. It does not contain any recognizable domains and a secondary structure prediction using Phyre2 does not give any clues as to its potential function (28). CD0580 (GapN) is annotated as an glyceraldehyde-3-phosphate dehydrogenase (GADPH), a key glycolytic enzyme, and contains an aldehyde dehydrogenase domain. Interestingly, its activity has been shown to be redox-controlled in other bacteria and has been implicated in the response to reactive oxygen and nitrogen species (29-31). CD1623 is a putative oxidoreductase with similarity to FAD flavoproteins and rubredoxins. CD1690 (TrxA) and CD1691 (TrxB) are likely encoded in the same operon (32), and form a thioredoxin/thioredoxin-disulfide reductase couple. CD0174 (CooS; IPR010047), CD0175 and CD0176 are likely also encoded in a single operon (32), and function as carbon monoxide dehydrogenase, and two putative oxidoreductases. As mentioned above, CD0174 has been implicated in aerobic/nitrosative stress and it is likely that CD1623, CD1690 and CD1691 also function in this pathway. Finally, CD2115A encodes another small hypothetical protein and as for CD0051A, no function could be assigned on the basis of secondary structure prediction.

As the σ^B^ regulon that we define here is substantially smaller than that previously reported, the major conclusion is that at least 32% of the σ^B^ regulon is involved in positively regulating oxidative/nitrosative stress responses. In the previous investigation of the σ^B^ regulon they were approximately 3.2% (∼32/1000) (19). Overall, we conclude that the core functions of the σ^B^ regulon lie in the regulation of the detoxification response to oxygen- and nitro-radicals.

### *In vitro* run-off transcriptions demonstrate direction activation of P_*cd0350*_, P_*cd2963*_, P_*cd3412*_ and P_*cd3605*_ by σ^B^

Gene expression can directly or indirectly be influenced by σ^B^, and to date no attempts have been made to discriminate these possibilities biochemically (11, 19). Despite the short time of induction and the uncoupling of σ^B^ from its normal regulatory network, our analyses could possibly also have picked up indirect effects. To determine if the transcription of selected genes is directly activated by σ^B^, *in vitro* transcription run-off reactions were performed using purified σ^B^_6xHis_ and RNA polymerase core enzyme (RNAP^core^) on the upstream regions of a selection of genes. The genes *cd0350* (encoding a putative hydrolase involved in oxidative stress), *cd2963* (encoding an L,D-transpeptidase), *cd3412* (encoding UvrB, involved in nucleotide excision repair), *cd3605* (encoding a ferredoxin) and *cd3614* (encoding a hypothetical protein involved in oxidative stress) were selected on the basis of our transcriptome analyses (**Dataset S1**), previous reporter gene assays (20) and/or the presence of a putative σ^B^ recognition site (11). The gene *cd0872* (maltose O-acetyltransferase) was not differentially expressed in our transcriptome data and was thus included as a negative control. The promoter of the toxin A gene (*tcdA*) in combination with purified TcdR was used as a positive control for the assay, as previously described (33).

As expected, no *in vitro* transcript was observed for a linear DNA fragment containing P_*cd0872*_ incubated with purified 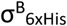and RNAP^core^ under our experimental conditions, whereas a specific product was obtained for the positive control P_*tcdA*_, in the presence of TcdR and RNAP^core^ (**Figure 4**). An RNAP^core^- and 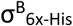-specific signal was observed for fragments containing the putative promoter regions of the genes *cd0350, cd2963, cd3412* and *cd3605*, demonstrating that expression of these genes was directed by σ^B^. For the fragments containing the putative promoter of *cd3614* we did not get a consistent product in the *in vitro* transcription experiments, though some smearing is visible in the lane with RNAP^core^ and 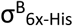. As *cd3614* demonstrates clear differential expression in the DNA array experiments and it upstream region harbors the σ^B^ consensus sequence WGWTT-N_13-17_-(G/T)GGTWA (19), we consider it likely that this gene is directly regulated by σ^B^ and our failure to obtain a discrete signal is due to our experimental conditions or the lack of an auxiliary factor in our *in vitro* assays.

**Figure 4.**
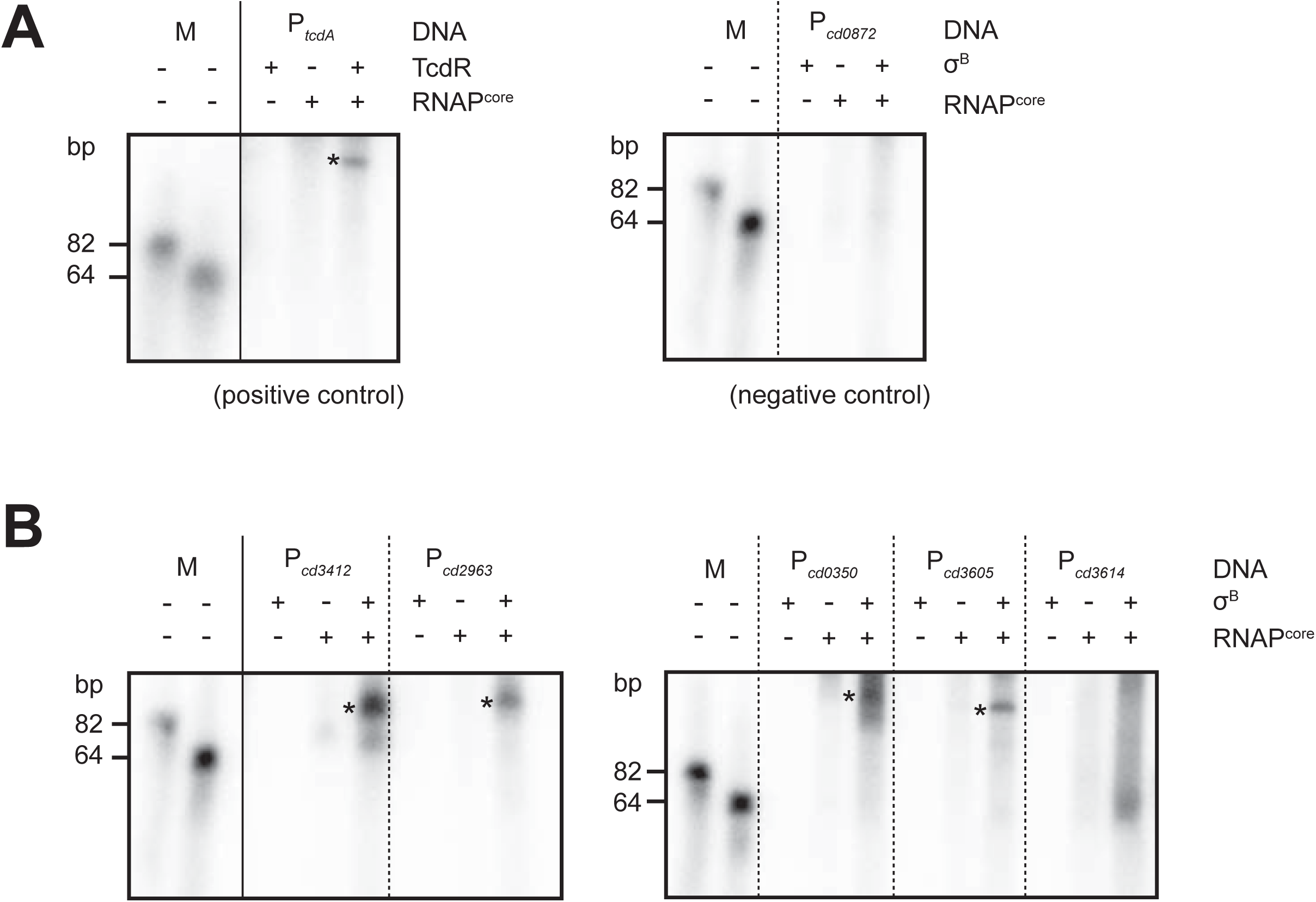
*In vitro* run-off transcription of selected promoter regions. Samples were run on an 8% urea gel. The two bands corresponding to 82 bp and 64 bp are end-labelled oligonucleotides. Reactions without sigma factor (σ^B^ or TcdR, respectively) or RNAP^core^ were analyzed as controls. Asterisks indicate the presence of distinct transcripts. RNAP^core^= *E. coli* RNA polymerase core enzyme (NEB). **A**) Controls for the *in vitro* run-off transcriptions. Purified TcdR, a sigma factor demonstrated to activate *tcdA* transcription *in vitro* (33), was used with P_*tcdA*_ (pCD22) as a positive control for the assay. P_*cd0872*_ (derived from pIB21) shows no altered transcription in the DNA array analysis and was taken along as negative control. **B**) *In vitro* run-off transcriptions for selected genes induced by σ^B^ overproduction.

Overall, we provide the first biochemical evidence for direct σ^B^-dependent activation of several genes identified via transcriptome analyses as part of the σ^B^ regulon in *C. difficile*.

### Antimicrobials and hydrogen peroxide activate σ^B^ -directed gene transcription

The redefined σ^B^ regulon pointed towards a substantial role for σ^B^ in coordinating the oxidative- and nitrosative stress response, which could result from antimicrobial treatment. In order to test for the activation of σ^B^-dependent promoters by antimicrobials, we set up a plate-based luciferase reporter assay. In this assay cells harboring σ^B^-dependent luciferase reporter constructs were plated on BHIY agar to give confluent growth and exposed to antimicrobials either through an epsilometer test (E-test) or through a filter disc. Subsequently, luciferase activity was imaged (for details, see Materials and Methods). A strain harboring P_tet_-*sluc*^*opt*^ (AP34) served as negative control, as this promoter is not expected to respond in a σ^B^-dependent manner (**Figure 5A**).

**Figure 5.**
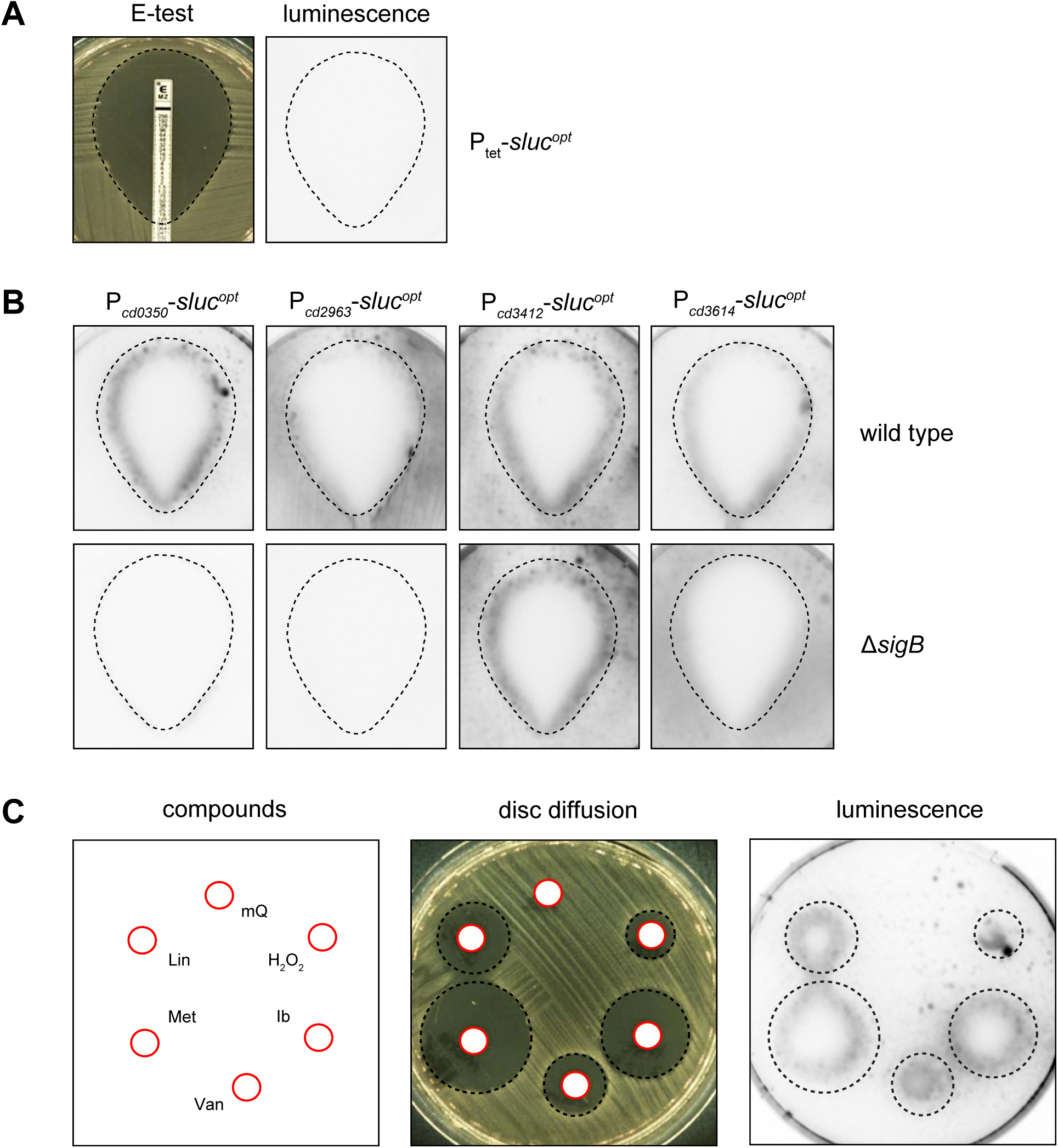
Plate-based luciferase assay shows σ^B^-dependant promotor activity from antimicrobial and hydrogen peroxide exposure. **A**) Setup of the assay. E-test halos (left panel) were sprayed with luciferase substrate and imaged (right panel). The dotted line indicates the location of the halo based on the left panel. Strain AP34 (P_tet_-*sluc*^opt^) shows no signal due to the absence of inducer. **B**) Luciferase reporters of different σ^B^ regulated promoters were tested for luciferase signal after a metronidazole E-test. Halos are indicated by the dashed lines as in panel A. **C**) Luciferase activity of the P_*cd0350*_ luciferase reporter was imaged in the presence of discs containing 10 μL of various compounds: sterile H_2_O (mQ), Lincomycin (3000 µg/ml; Lin), metronidazole (200 µg/ml; Met), vancomycin (200 µg/ml; Van), Ibezapolstat (400 µg/ml; Ib) and hydrogen peroxide (1M; H_2_O_2_). Halos and the location of the different stressors are indicated by red circles (disc) and black dashed circles (halo).

First, the σ^B^-dependent response to metronidazole was investigated. Metronidazole, formerly used as a first line treatment for CDI, is believed to cause DNA damage through the formation of nitro-radicals although its exact mode of action remains unclear (6, 34). To survey a full spectrum of metronidazole concentrations, we evaluated luminescence after 24h of incubation of an metronidazole E-test. If metronidazole treatment results in σ^B^-dependent activation of gene transcription, we expect to see a luciferase signal in the wild type, but not in a σ^B^ knockout background. In agreement with this, activation of P_*cd0350*_ was observed at the edge of the halo resulting from the metronidazole E-test in the wild type background, but not in the σ^B^ knockout strain (**Figure 5B**). No signal was observed for the negative control P_tet_-*sluc*^*opt*^ (**Figure 5A**). The observed activation σ^B^-dependent activation of gene expression at the edge of the halo, but not further into the plate suggests that the metronidazole induced, σ^B^-dependent activation of P_*cd0350*_ occurred close to the Minimal Inhibitory Concentration (MIC). Expression of the luciferase from P_*cd296*3_ was found to be strictly dependent on σ^B^ as no luciferase activity was observed in the *sigB* knockout strain. However, there was limited to no increase in reporter gene expression in the presence of metronidazole. Metronidazole strongly activated transcription from P_*cd3412*_ at MIC levels of metronidazole, but this appeared to be independent of σ^B^ in this assay since in the *sigB* mutant a similar induction was observed. Finally, in a manner comparable to P_*cdD0350*_, the activation of P_CD3614_ was strongly induced by metronidazole at values close to MIC in a σ^B^-dependent manner, but residual activity was observed in the σ^B^ knockout strain independent of metronidazole levels. We noted that metronidazole-induced promoter activation appeared to occur on the inside of the E-test halo, which might be attributed to the secretion of the luciferase reporter.

The observed diverse regulatory responses at different tested promoters during the treatment of *C. difficile* with metronidazole pointed towards a more complex regulatory network with the participation of σ^B^. Antimicrobial-driven (and σ^B^-dependent) activation of σ^B^ target genes could be specific to metronidazole or represent a more general response to cellular (toxic) stresses. Therefore, we evaluated the effects of different antimicrobial compounds and the radical producer H_2_O_2_ as a positive control (19), using the P_*cd0350*_ reporter construct, as this promoter demonstrated the clearest σ^B^ dependent activation in the presence of metronidazole (**Figure 5B**). We tested the cell wall biosynthesis inhibitor vancomycin, the protein synthesis inhibitor lincomycin and the DNA polymerase inhibitor ibezapolstat (formerly known as ACX-362E) (35). We observed clear activation of P_*cd0350*_ in the presence of all added stressors, but not for a negative control containing water (**Figure 5C**).

We conclude that, at least for the σ^B^ dependent promoter of *cd0350*, activation does not only occur upon exposure to lethal levels of metronidazole, but also with unrelated antimicrobials and toxic stressors such as hydrogen peroxide.

## Discussion

In this work we have demonstrated by Western blotting using an affinity purified anti-σ^B^ antibody that σ^B^ can be overproduced for a limited period of time, sufficient for transcriptome analyses. The induced production of σ^B^ in a *sigB* mutant background yielded highly consistent results, despite potential toxicity and plasmid-loss (**Figure 2**), and the results were used to redefine the surprisingly large σ^B^ regulon previously proposed (19). As our approach more accurately measures changes in transcription directly related to σ^B^ production, the refined regulon described here is much smaller (**Dataset S1**). Its size is fully in line with that of the σ^B^ regulon of other gram-positive bacteria such as *L. monocytogenes* (∼130 genes), *B. subtilis* (∼150 genes) and *S. aureus* (∼200 genes) (8). The redefined regulon underscores the importance of σ^B^ in responding to oxidative/nitrosative stresses as genes implicated in such processes are significantly enriched in the smaller regulon.

The majority of the genes in our regulon were found to be induced, rather than repressed, by σ^B^. This is in line with sigma factors acting as specificity determinants for transcription initiation (36). Similar observations have been made for the σ^B^ regulon of *L. monocytogenes* (37, 38). For the first time, direct evidence of *C. difficile* σ^B^-dependent gene activation is provided by the results of the *in vitro* run-off transcriptions (**Figure 4**), which demonstrate that RNAP^core^ and σ^B^ are sufficient to generate transcripts from P*cd0350*, P*cd2963*, P*cd3412* and P*cd3605*. Notably, these experiments pave the way for a further *in vitro* characterization of this sigma factor in *C. difficile* including validation of the σ^B^ binding sequence and the interplay with other regulators.

Although the promoters of *cd3412* (*uvrB*) and *cd3614* were reported to have a σ^B^ consensus sequence and are differentially expressed upon σ^B^ overexpression (19, 35), our results clearly demonstrate that they can also be expressed in a σ^B^-independent manner (**Figure 5B**). This is most notable for P_*cd3412*_, which is still activated by metronidazole in the absence of σ^B^, in line with results obtained with ibezapolstat in a different study (20). Both metronidazole and ibezapolstat treatment can cause DNA damage, and DNA-damage dependent induction of *cd3412* therefore likely depends on a *sigB*-independent pathway. The observed σ^B^-dependent gene repression could be indirect (σ^B^ induces the transcription of a repressor gene), or the result of competition (σ^B^ competes with other sigma factors for RNAP). We consider the second scenario more likely for the following reasons. First, little overproduction of σ^B^ was detected after 30 minutes of induction. This leaves only a limited time for indirect effects to occur in our setup. Second, the majority of genes down-regulated upon overexpression of σ^B^ fall into a single functional group (flagellar motility). These genes are known to be regulated by the dedicated sigma factor, σ^D^ (39), supporting the model of sigma factor competition. Strikingly, in *L. monocytogenes* σ^B^ activity (indirectly) also results in down-regulation of flagellar gene expression, but this is mediated by the repressor MogR (40). Protein BLAST analyses revealed that *C. difficile* does not possess a MogR homologue. Nevertheless, the conserved inverse correlation between the σ^B^-dependent general stress response and bacterial motility could represent a cost-saving strategy for bacterial cells (41). The indirect mechanism underlying the observed σ^B^-dependent down-regulation in *C. difficile* remains to be determined.

There appears to be an intriguing link between σ^B^ and the response to toxic compounds, as a *sigB* mutant was more susceptible to rifampicin and mitomycin C (19) and exposure to antimicrobials (metronidazole, vancomycin, lincomycin and ibezapolstat) or hydrogen peroxide leads to σ^B^-dependent promoter activation (**Figure 5C**). The mechanism behind the latter is unclear. It has been suggested that antimicrobials at toxic concentrations can influence metabolism and respiration (42, 43), potentially resulting in the formation of bactericidal concentrations of radical species (44-46). A strong connection between σ^B^ and oxidative (and/or nitrosative) stress in *C. difficile* (**Table 2**) and other bacteria (7, 18, 19), as well as a recently described radical scavenging strategy that increases tolerance to antimicrobials (47) are consistent with such a model. However, additional research is necessary to determine exactly how these processes occur and are influenced by antimicrobials in anaerobic organisms under anoxic conditions.

In conclusion, we have demonstrated that σ^B^ is directly involved in metabolic and oxidative stress responses and that lethal stresses may influence these processes, resulting in activation of σ^B^ targeted genes.

## Acknowledgements

We would like to thank Bruno Dupuy for kindly providing us with plasmid pCD22 and Annemieke Friggen for technical assistance in strain construction. We thank Nicholas Kint and Isabelle Martin-Verstraete for communicating results prior to publication and Joachim Goedhart for advice in using VolcanoseR.

## Materials and Methods

### Construction of σ^B^ expression- and luciferase-reporter vectors

All oligonucleotides used in this study can be found in table 2. Plasmids and strains can be found in table 3. The P_T7_-*sigB*_6x_ expression vector pIB14 was created by restriction-ligation using restriction enzymes NdeI and XhoI. Using primers oIB-1 and oIB-2 on *C. difficile* 630Δ*erm* chromosomal DNA, the *sigB* CDS was amplified by PCR. All PCR products used for sequencing or plasmid synthesis were performed with Q5 high-fidelity polymerase (NEB). The resulting DNA fragment was digested and ligated into NdeI-xhoI digested pET21b(-) vector, generating expression vector pIB14. Plasmids pIB27, pIB68, pIB69 and pIB74 have been described previously (20). The CD0872 promoter area was amplified using primers oIB-14 and oIB15 and the P_*cd0872*_ luciferase-reporter plasmid was created by restriction-ligation using restriction enzymes KpnI and SacI in digested pAP24 backbone, generating plasmid pIB21. Using Gibson assembly as described in (20) the P_*cd3605*_ luciferase-reporter plasmid was generated using primers oIB-90 and oIB-99, yielding plasmid pIB73. A plasmid containing P_tet_-*sigB* was generated by cloning the 630Δ*erm sigB* CDS amplified with oWKS-1498 and oWKS-1499 in pMiniT (NEB E1202) per manufacturer’s instructions. Using restriction enzymes SacI and BamHI this PCR fragment was cloned into pRPF185 generating pWKS1760. All plasmids were verified by Sanger sequencing.

**Table 3.**
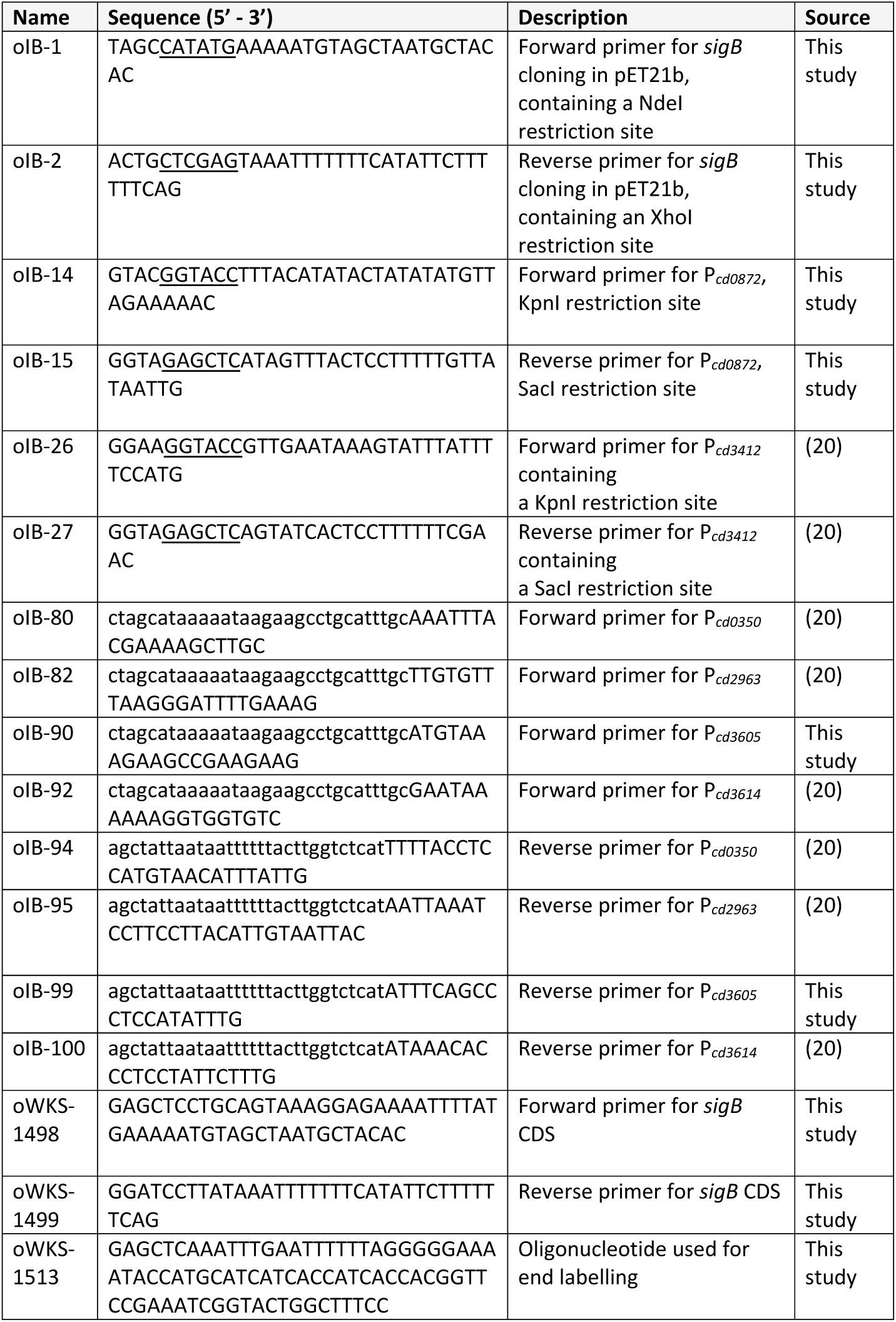
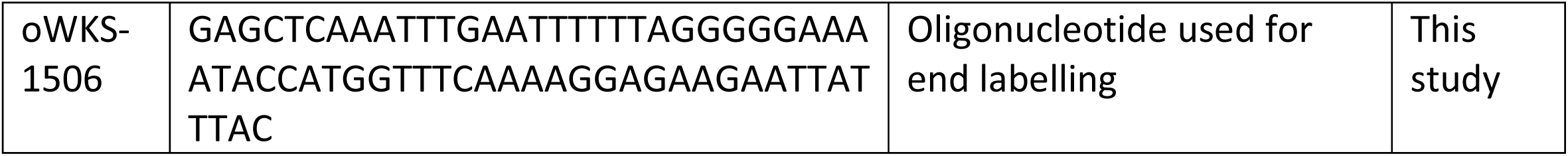
Oligonucleotides used in this study. Restriction sites are underlined, 30 bp overlapping regions as used in Gibson Assembly are indicated in lowercase letters.

**Table 4.** Plasmids and strains used in this study.

|  | Name | Relevant features | Source |
| --- | --- | --- | --- |
| Plasmids | pAP24 | P<sub>tet</sub>-*sluc*<sup>opt</sup>; *catP* | (24) |
|  | pCD22 | P<sub>tcdA</sub>; *catP* | (53) |
|  | pIB14 | P<sub>T7</sub>-*sigB*<sub>6xHis</sub>; *amp* | This study |
|  | pIB21 | P<sub>cd0872</sub>-*sluc*<sup>opt</sup>; *catP* | This study |
|  | pIB27 | P<sub>cd3412</sub>-*sluc*<sup>opt</sup>; *catP* | (20) |
|  | pIB68 | P<sub>cd0350</sub>-*sluc*<sup>opt</sup>; *catP* | (20) |
|  | pIB69 | P<sub>cd2963</sub>-*sluc*<sup>opt</sup>; *catP* | (20) |
|  | pIB73 | P<sub>cd3605</sub>-*sluc*<sup>opt</sup>; *catP* | This study |
|  | pIB74 | P<sub>cd3614</sub>-*sluc*<sup>opt</sup>; *catP* | (20) |
|  | pWKS1750 | P<sub>tet</sub>-*sigB*; *catP* | This study |
| Strains | 630Δ*erm* | MLS-susceptible derivative of *Clostridioides difficile* strain 630 | (54) or (55) |
| | AP34 | 630Δ*erm* pAP24; thia<sup>R</sup> | (24) |
| | DH5α | *Escherichia coli* F<sup>-</sup> *endA1 glnV44 thi-1 recA1 relA1 gyrA96 deoR nupG purB20 ϕ80dlacZΔM15 $\Delta$(lacZYA-argF)U169 hsdR17(r<sub>K</sub><sup>-</sup> m<sub>K</sub><sup>+</sup>) λ<sup>-</sup>* | Laboratory stock |
|  | IB14 | Rosetta (DE3) pLysS pIB14; amp<sup>R</sup>, chlor<sup>R</sup> | This study |
| | IB18 | 630Δ*erm* *sigB::CT*; erm<sup>R</sup>/linco<sup>R</sup> | This study |
| | IB37 | 630Δ*erm* pIB27; thia<sup>R</sup> | (20) |
| | IB56 | 630Δ*erm* Δ*sigB* | (20) |
|  | IB58 | IB18 pWKS1750; thia<sup>R</sup>, erm<sup>R</sup>/linco<sup>R</sup> | This study |
|  | IB61 | IB18 pAP24; thia<sup>R</sup>, erm<sup>R</sup>/linco<sup>R</sup> | This study |
| | IB95 | 630Δ*erm* pIB68; thia<sup>R</sup> | (20) |
| | IB96 | 630Δ*erm* pIB69; thia<sup>R</sup> | (20) |
|  | IB98 | IB56 pIB27; thia<sup>R</sup> | (20) |
|  | IB99 | IB56 pIB68; thia<sup>R</sup> | (20) |
|  | IB100 | IB56 pIB69; thia<sup>R</sup> | (20) |
| | IB108 | 630Δ*erm* pIB74; thia<sup>R</sup> | (20) |
|  | IB111 | IB56 pIB74; thia<sup>R</sup> | (20) |
| | JC096 | 630Δ*erm* pWKS1750; thia<sup>R</sup> | This study |

### Bacterial strains and growth conditions

Strains of *E. coli* were grown aerobically at 37°C in Luria-Bertani broth (Affymetrix) supplemented with ampicillin (50 ug/mL), kanamycin (50 ug/mL) and/or chloramphenicol (20 ug/mL) when required. Plasmids were maintained in *E. coli* strains DH5α or MDS42 (Scarab Genomics) under appropriate antimicrobial selection and cells were transformed using standard procedures (48). For plasmid conjugation into recipient *C. difficile* 630*Δerm, E. coli* strain CA434 was used as a donor strain as previously described (49). *C. difficile* strains were cultured anaerobically at 37°C in either a Don Whitley VA-1000 or A55 workstation. Cells were cultured in Brain Heart Infusion (BHI, Oxoid) broth supplemented with 0.5% (w/v) yeast-extract (BHIY) and 20 µg/ml thiamphenicol when appropriate. Unless additional antimicrobials/stressors were added (metronidazole E-test and sterile pads supplemented with different stressors), medium was supplemented with *C. difficile* Selective Supplements (CDSS, Oxoid).

### Overproduction, purification and affinity purification of 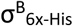 for synthesis of a polyclonal anti-σ^B^ antibody

#### Overproduction and purification of 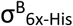

Overexpression of 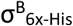 was performed by using Escherichia coli (*E. coli*) Rosetta (DE3) pLysS cells (Novagen) harboring the *E. coli* expression plasmid pIB14. These cells were cultured in Luria-Bertani (LB) broth and induced with 0.5 mM IPTG for one hour starting at an optical density ≈ 0.6. Cells were collected by centrifugation at 4°C and pellets were resuspended in lysis buffer (pH=8.0, 50 mM NaH_2_PO_4_, 300 mM NaCl, 5mM β-mercaptoethanol, 0.1% NP-40 and Complete protease inhibitor cocktail (CPIC, Roche Applied Science). Through the addition of 1 mg/mL lysozyme and sonication (6 times 20 seconds) cells were lysed. The lysate was drawn through a blunt 1.2mm needle and was clarified by centrifugation at 13000 x*g* at 4°C for 25 minutes. The supernatant containing recombinant 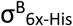 was purified on TALON Superflow resin (GE healthcare) per manufacturer’s instructions. Proteins were dialyzed and stored in buffer (pH=8.0) containing 50 mM NaH_2_PO_4_, 300 mM NaCl and 12% glycerol. Protein concentrations were determined using a Bradford assay (Biorad) and 2 mL of 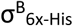 protein solution containing 2 mg/mL measured protein was sent to BioGenes GmbH (Berlin) for generation of a polyclonal rabbit-anti-σ^B^ antibody.

#### Affinity purification of the polyclonal anti-σ^B^ antibody

Affinity purification of the antibody was performed to increase specificity. Approximately 350µg of purified 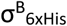 protein was loaded on an SDS-PAGE gel. After transferring proteins to a PVDF membrane using standard blotting procedures, purified 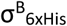 protein was visualized by Ponceau S stain and the membrane containing the protein was cut as small as possible whilst retaining the region with the protein. The membrane was destained and washed with TBST buffer (500 mM NaCl, 20 mM Tris-base, 0.05% v/v Tween-20, pH=7.4) twice for 5 minutes at room temperature. The membrane was then pre-eluted by soaking in acidic glycine solution (100 mM, pH=2.5) for 5 minutes prior to washing with TBST twice for five minutes at room temperature. Subsequently the membrane was blocked in 5% non-fat milk powder solution (Campina Elk, dissolved in TBST buffer) for one hour at room temperature after again washing twice with TBST for 5 minutes. Serum containing anti-σ^B^ antibody was incubated on the membrane overnight at 4°C. After three 5-minute washes with TBST, the membrane was washed twice for 5 minutes in PBS. Affinity purified antibody was eluted from the membrane by adding acidic glycine solution and incubating for 10 minutes at room temperature. The pH of the eluate was adjusted to 7.0 through the addition of 1M Tris-HCl (pH=8.0). This step was repeated twice more and the eluates were pooled and centrifuged (1 min maximum speed) to remove precipitated protein and membrane particles. Bovine serum albumin (BSA) and sodium azide were added to the affinity purified anti-σ^B^ antibody to end concentrations of 1 mg/mL and 5mM respectively, and the affinity purified antibody was stored at -80°C.

### Characterization of the σ^B^ regulon

#### *σ*^*B*^ *overproduction in* C. difficile

Exponentially growing starter cultures of IB58 and IB61 were diluted to an OD_600_=0.05 in BHIY supplemented with 20 µg/mL lincomycin with thiamphenicol (20 µg/mL) where appropriate. Cells were grown until OD_600_≈0.3 after which 1 mL sample was taken for control by Western blot and cells were induced with 100 ng/mL ATc for one hour. After one hour of induction, 1 mL of sample was taken for control by Western Blot and 50 mL was collected by centrifugation and stored at -20°C until RNA extraction. Non-induced samples were treated and collected identically, except that no ATc was added at OD_600_≈0.3. All samples were corrected for OD_600_ prior to analysis by Western Blot.

#### RNA extraction

Bacterial RNA was extracted and analyzed as previously described (50). Briefly, cell pellets were lysed in enzymatic lysis buffer consisting of 15 mg/mL lysozyme and TE buffer. Further disruption of cells was performed by vigorous mechanical lysis for 3 minutes in RLT buffer to which one spatula of glass beads was added. After samples were centrifuged (3 minutes at 10.000 rpm at 4°C) and 100% ethanol was added to the supernatant, RNA was purified using the Qiagen RNeasy Kit protocol according to manufacturer’s instructions. DNA contamination was removed by using RNAse-free DNAse I (Qiagen) twice prior to elution of the RNA samples in H_2_O. RNA quality and integrity numbers (RINs) were assessed with a Bioanalyzer 2100 (Agilent) and RNA 6000 Nano Reagents (Agilent). Only samples with an RIN of≥ 7 were used for analysis.

#### DNA microarray and data analysis

A customized whole-genome DNA microarray of 630Δ*erm* was used (8×15K format, Agilent) (50). Quadruplicate samples were analyzed for the DNA microarray. Using the ULS fluorescent labelling kit for Agilent arrays (Kreatech), 1 µg of total RNA was used for labelling with either Cy3 (P_tet_-*sigB*) or Cy5 (P_tet_-*sluc*^*opt*^). After pooling and fragmentation, 300 ng of labelled RNA per sample was hybridized according to the two-color microarray protocol from Agilent. DNA microarrays were scanned with an Agilent C scanner and analyzed as described (50). A gene was considered differentially expressed if the log2 foldchange (log2 FC) was ≤ -1.5 or ≥1.5 and the p-value was <0.05. Results were visualized in VolcanoseR (27), and are available as an interactive graph via the URL contained in **Text S1**. The data used in the visualization has been deposited at Zenodo for this purpose (doi: 10.5281/zenodo.3945936). Full transcriptome data has been deposited in the GEO database, and can be accessed through the identifier GSE152515.

### *In vitro* transcription

DNA oligonucleotides oWKS-1506 and oWKS-15136 (64- and 82 bp, respectively) were end-labelled with ÿ-32^P^-ATP using T4 Polynucleotide Kinase (PNK) and used as a size indicator for the *in vitro* transcription reactions. For the end labelling reaction, 1 µl ÿ-32P-ATP was incubated together with 200 pmol oligonucleotide and 1 µl PNK in Forward Reaction buffer (70 mM Tris-HCl (pH 7.6), 10 mM MgCl_2_, 100 mM KCl, 1 mM 2-mercaptoethanol) at 37oC for 30 minutes. For the *in vitro* run-off transcriptions sigma factors and RNA polymerase core enzyme were preincubated with PCR amplified promoter areas (for P_*CD0350*_, P_*CD0872*_, P_*CD2963*_, P_*CD3412*,_ P_*CD3605*_ and P_*CD3614*_) or XbaI linearised pCD22 (P_*tcdA*_) for 30 minutes at 37oC prior to start of the reaction. PCR products of the promoter areas as used for the *in vitro* transcription reactions were loaded on and excised from agarose gels and purified using a NucleoSpin™ Gel and PCR Clean-up kit (Machery-Nagel). *In vitro* transcription reactions mixtures contained 1 µl *E. coli* RNAP^core^ (NEB M0550S)), 16 pmol sigma factor, 0.5 pmol DNA, 10 mM NTP mix and 0.3 µl α-32^P^-ATP in reaction buffer (40mM Tris-HCl, 150 mM KCl, 10 mM MgCl_2_, 1mM DTT, 0.01% triton X-100, pH=7.5) and were incubated for 15 minutes at 37°C. Transcripts and labelled oligonucleotides to be used as a size indication were purified using P-30 Bio-Gel spin columns (Biorad). All reactions were stopped in gel loading buffer II (Invitrogen) containing 95% Formamide, 18 mM EDTA and 0.025% each of SDS, xylene cyanol, and bromophenol blue at 95°C for 5 minutes and loaded on 8% monomeric UreaGel (SequaGel, National Diagnostics). Gels were dried and exposed to PhosphorImager screens overnight (approximately 17 hours) and imaged with a Typhoon-9410 scanner (GE Healthcare).

### Spot assay for viability upon σ^B^ overproduction in *C. difficile* and vector stability assay

*C. difficile* overnight pre-cultures were corrected for OD_600_ and subsequently tenfold serially diluted in BHI medium. Two-µl spots of each dilution were plated on selective (20 µg/mL thiamphenicol) and unselective square (90×15 mm, VWR international) BHI plates with or without 200 ng/mL anhydrotetracycline (ATc). Growth was evaluated after 24 hours and subsequently swabs were taken from all strains grown on unselective BHI agar plates with- and without 200 ng/mL ATc for the vector stability assay. These swabs used for the vector stability assay were resuspended in BHI medium, adjusted for OD_600_-values and tenfold serially diluted in unselective BHI medium. Of these OD-corrected, serially diluted suspensions, 10-µl spots were then plated on selective (20 µg/mL thiamphenicol + CDSS) and non-selective (BHI +CDSS) plates and CFU/mL was counted after 24-48 hours of growth. Percentage plasmid maintained was calculated as [CFU/mL]_selective_/[CFU/mL]_non-selective_ x 100%If no growth was detected on selective plates percentage of plasmid maintained was set as 0% To calculate statistical significance between % plasmid maintained in strains induced- or not induced by ATc, an unpaired student’s t-test was used.

### Plate based luciferase assay with metronidazole E-test and disk diffusion

Strains harboring luciferase-reporter plasmids were grown on prereduced, selective BHI plates for 24 hours. Subsequently bacterial suspensions corresponding to 1.0 McFarland turbidity were applied on BHI agar supplemented with 0.5% yeast-extract after which a metronidazole E-test or plain disks were applied. Disks were spotted with 10 µl each of sterile H_2_O, 1M H_2_O_2_, 3000 µg/mL lincomycin, 200 µg/mL metronidazole, 400 µg/mL ibezapolstat and 200 µg/mL vancomycin. After 24 hours of growth luciferase activity was visualized by spraying one spray of 1:100 reconstituted NanoGlo Luciferase substrate (Promega N1110) per agar plate using a disposable spray flask. One spray corresponded to approx. 250 µl reconstituted NanoGlo Luciferase substrate. Luminescence was recorded using a Uvitec Alliance Q9 Advanced imager (BioSPX) after 10 seconds exposure time per plate. Reporter constructs were generated in a *sigB* knockout made by Allelic Coupled Exchange (51) as opposed to the ClosTron mutant background used for the DNA arrays. However, no differences between these backgrounds has ever been observed in our assays. on factor

## Legends for Supplemental Material

**Dataset S1**. Genes differentially expressed in σ^B^ overproducing cells compared to controls. Cells were harvested from cultures with (induced) or without (uninduced) ATc, and with (+Tm) or without (-Tm) thiamphenicol. Gene name = generic gene name (or locus tag if not available). log2FC is the log2 of the fold change in gene expression. Four different comparisons are shown, genes are aligned between comparisons. Up-regulated genes are indicated in green. Down-regulated genes are indicated in red. Genes not considered not part of the σ^B^ regulon are highlighted in yellow.

**Dataset S2**. Manhattan distance for differentially regulated genes from hybridization 2 (cells harvested from cultures grown in the presence of both ATc and thiamphenicol). Change indicates whether expression is higher (increased) or lower (decreased) upon overproduction of σ^B^. log2FC = log2 of the fold change in gene expression. Significance = -10log of the adjusted p-value (see Dataset S1). Manhattan distance was calculated by and exported using the online tool VolcanoseR.

**Text S1**. Link to access an interactive version of the volcano plot based on hybridization 2 (cells harvested from cultures grown in the presence of both anhydrotetracyclin and thiamphenicol).

## References

1. Chang JY, Antonopoulos DA, Kalra A, Tonelli A, Khalife WT, Schmidt TM, Young VB. 2008. Decreased diversity of the fecal Microbiome in recurrent Clostridium difficile-associated diarrhea. J Infect Dis 197:435–8.

2. Smits WK, Lyras D, Lacy DB, Wilcox MH, Kuijper EJ. 2016. Clostridium difficile infection. Nat Rev Dis Primers 2:16020.

3. Paredes-Sabja D, Shen A, Sorg JA. 2014. Clostridium difficile spore biology: sporulation, germination, and spore structural proteins. Trends Microbiol 22:406–16.

4. Fang FC, Frawley ER, Tapscott T, Vazquez-Torres A. 2016. Bacterial Stress Responses during Host Infection. Cell Host Microbe 20:133–43.

5. McDonald LC, Gerding DN, Johnson S, Bakken JS, Carroll KC, Coffin SE, Dubberke ER, Garey KW, Gould CV, Kelly C, Loo V, Shaklee Sammons J, Sandora TJ, Wilcox MH. 2018. Clinical Practice Guidelines for Clostridium difficile Infection in Adults and Children: 2017 Update by the Infectious Diseases Society of America (IDSA) and Society for Healthcare Epidemiology of America (SHEA). Clin Infect Dis 66:987–994.

6. Abt MC, McKenney PT, Pamer EG. 2016. Clostridium difficile colitis: pathogenesis and host defence. Nat Rev Microbiol 14:609–20.

7. Emerson JE, Stabler RA, Wren BW, Fairweather NF. 2008. Microarray analysis of the transcriptional responses of Clostridium difficile to environmental and antibiotic stress. J Med Microbiol 57:757–64.

8. Guldimann C, Boor KJ, Wiedmann M, Guariglia-Oropeza V. 2016. Resilience in the Face of Uncertainty: Sigma Factor B Fine-Tunes Gene Expression To Support Homeostasis in Gram-Positive Bacteria. Appl Environ Microbiol 82:4456–69.

9. van Schaik W, Abee T. 2005. The role of sigmaB in the stress response of Gram-positive bacteria -- targets for food preservation and safety. Curr Opin Biotechnol 16:218–24.

10. van Schaik W, Tempelaars MH, Zwietering MH, de Vos WM, Abee T. 2005. Analysis of the role of RsbV, RsbW, and RsbY in regulating {sigma}B activity in Bacillus cereus. J Bacteriol 187:5846–51.

11. Kint N, Alves Feliciano C, Hamiot A, Denic M, Dupuy B, Martin-Verstraete I. 2019. The sigma(B) signalling activation pathway in the enteropathogen Clostridioides difficile. Environ Microbiol 21:2852–2870.

12. Guerreiro DN, Wu J, Dessaux C, Oliveira AH, Tiensuu T, Gudynaite D, Marinho CM, Boyd A, García-del Portillo F, Johansson J, O’Byrne CP. 2020. Mild stress conditions during laboratory culture promote the proliferation of mutations that negatively affect Sigma B activity in <em>Listeria monocytogenes</em>. Journal of Bacteriology doi: 10.1128/jb.00751-19:JB.00751-19.

13. Boylan SA, Thomas MD, Price CW. 1991. Genetic method to identify regulons controlled by nonessential elements: isolation of a gene dependent on alternate transcription factor sigma B of Bacillus subtilis. J Bacteriol 173:7856–66.

14. Garner MR, James KE, Callahan MC, Wiedmann M, Boor KJ. 2006. Exposure to salt and organic acids increases the ability of Listeria monocytogenes to invade Caco-2 cells but decreases its ability to survive gastric stress. Appl Environ Microbiol 72:5384–95.

15. Kim H, Boor KJ, Marquis H. 2004. Listeria monocytogenes sigmaB contributes to invasion of human intestinal epithelial cells. Infect Immun 72:7374–8.

16. Morikawa K, Maruyama A, Inose Y, Higashide M, Hayashi H, Ohta T. 2001. Overexpression of sigma factor, sigma(B), urges Staphylococcus aureus to thicken the cell wall and to resist beta-lactams. Biochem Biophys Res Commun 288:385–9.

17. DeMaio J, Zhang Y, Ko C, Bishai WR. 1997. Mycobacterium tuberculosis sigF is part of a gene cluster with similarities to the Bacillus subtilis sigF and sigB operons. Tuber Lung Dis 78:3–12.

18. Bandow JE, Brotz H, Hecker M. 2002. Bacillus subtilis tolerance of moderate concentrations of rifampin involves the sigma(B)-dependent general and multiple stress response. J Bacteriol 184:459–67.

19. Kint N, Janoir C, Monot M, Hoys S, Soutourina O, Dupuy B, Martin-Verstraete I. 2017. The alternative sigma factor sigma(B) plays a crucial role in adaptive strategies of Clostridium difficile during gut infection. Environ Microbiol 19:1933–1958.

20. van Eijk E, Boekhoud IM, Kuijper EJ, Bos-Sanders IMJG, Wright G, Smits WK. 2019. Genome Location Dictates the Transcriptional Response to PolC Inhibition in <em>Clostridium difficile</em>. 63:e01363–18.

21. Fagan RP, Fairweather NF. 2011. Clostridium difficile has two parallel and essential Sec secretion systems. J Biol Chem 286:27483–93.

22. Ransom EM, Ellermeier CD, Weiss DS. 2015. Use of mCherry Red fluorescent protein for studies of protein localization and gene expression in Clostridium difficile. Appl Environ Microbiol 81:1652–60.

23. Boekhoud IM, Hornung BVH, Sevilla E, Harmanus C, Bos-Sanders IMJG, Terveer EM, Bolea R, Corver J, Kuijper EJ, Smits WK. 2020. Plasmid-mediated metronidazole resistance in Clostridioides difficile. Nature Communications 11:598.

24. Oliveira Paiva AM, Friggen AH, Hossein-Javaheri S, Smits WK. 2016. The Signal Sequence of the Abundant Extracellular Metalloprotease PPEP-1 Can Be Used to Secrete Synthetic Reporter Proteins in Clostridium difficile. ACS Synth Biol 5:1376–1382.

25. Gao X, Dong X, Subramanian S, Matthews PM, Cooper CA, Kearns DB, Dann CE, 3rd. 2014. Engineering of Bacillus subtilis strains to allow rapid characterization of heterologous diguanylate cyclases and phosphodiesterases. Appl Environ Microbiol 80:6167–74.

26. Poquet I, Saujet L, Canette A, Monot M, Mihajlovic J, Ghigo JM, Soutourina O, Briandet R, Martin-Verstraete I, Dupuy B. 2018. Clostridium difficile Biofilm: Remodeling Metabolism and Cell Surface to Build a Sparse and Heterogeneously Aggregated Architecture. Front Microbiol 9:2084.

27. Goedhart J, Luijsterburg MS. 2020. VolcaNoseR – a web app for creating, exploring and sharing volcano plots. bioRxiv doi: https://doi.org/10.1101/2020.05.07.082263.

28. Kelley LA, Mezulis S, Yates CM, Wass MN, Sternberg MJ. 2015. The Phyre2 web portal for protein modeling, prediction and analysis. Nat Protoc 10:845–58.

29. Hillion M, Imber M, Pedre B, Bernhardt J, Saleh M, Loi VV, Maass S, Becher D, Astolfi Rosado L, Adrian L, Weise C, Hell R, Wirtz M, Messens J, Antelmann H. 2017. The glyceraldehyde-3-phosphate dehydrogenase GapDH of Corynebacterium diphtheriae is redox-controlled by protein S-mycothiolation under oxidative stress. Sci Rep 7:5020.

30. Christodoulou D, Link H, Fuhrer T, Kochanowski K, Gerosa L, Sauer U. 2018. Reserve Flux Capacity in the Pentose Phosphate Pathway Enables Escherichia coli’s Rapid Response to Oxidative Stress. Cell Syst 6:569–578 e7.

31. Imber M, Huyen NTT, Pietrzyk-Brzezinska AJ, Loi VV, Hillion M, Bernhardt J, Tharichen L, Kolsek K, Saleh M, Hamilton CJ, Adrian L, Grater F, Wahl MC, Antelmann H. 2018. Protein S-Bacillithiolation Functions in Thiol Protection and Redox Regulation of the Glyceraldehyde-3-Phosphate Dehydrogenase Gap in Staphylococcus aureus Under Hypochlorite Stress. Antioxid Redox Signal 28:410–430.

32. Dehal PS, Joachimiak MP, Price MN, Bates JT, Baumohl JK, Chivian D, Friedland GD, Huang KH, Keller K, Novichkov PS, Dubchak IL, Alm EJ, Arkin AP. 2010. MicrobesOnline: an integrated portal for comparative and functional genomics. Nucleic Acids Res 38:D396–400.

33. Matamouros S, England P, Dupuy B. 2007. Clostridium difficile toxin expression is inhibited by the novel regulator TcdC. Mol Microbiol 64:1274–88.

34. Edwards DI. 1993. Nitroimidazole drugs-action and resistance mechanisms I. Mechanism of action. Journal of Antimicrobial Chemotherapy 31:9–20.

35. van Eijk E, Boekhoud IM, Kuijper EJ, Bos-Sanders I, Wright G, Smits WK. 2019. Genome Location Dictates the Transcriptional Response to PolC Inhibition in Clostridium difficile. Antimicrob Agents Chemother 63.

36. Helmann JD. 2019. Where to begin? Sigma factors and the selectivity of transcription initiation in bacteria. Mol Microbiol 112:335–347.

37. Mujahid S, Orsi RH, Vangay P, Boor KJ, Wiedmann M. 2013. Refinement of the Listeria monocytogenes sigmaB regulon through quantitative proteomic analysis. Microbiology 159:1109–1119.

38. Palmer ME, Chaturongakul S, Wiedmann M, Boor KJ. 2011. The Listeria monocytogenes sigmaB regulon and its virulence-associated functions are inhibited by a small molecule. mBio 2.

39. El Meouche I, Peltier J, Monot M, Soutourina O, Pestel-Caron M, Dupuy B, Pons JL. 2013. Characterization of the SigD regulon of C. difficile and its positive control of toxin production through the regulation of tcdR. PLoS One 8:e83748.

40. Toledo-Arana A, Dussurget O, Nikitas G, Sesto N, Guet-Revillet H, Balestrino D, Loh E, Gripenland J, Tiensuu T, Vaitkevicius K, Barthelemy M, Vergassola M, Nahori MA, Soubigou G, Regnault B, Coppee JY, Lecuit M, Johansson J, Cossart P. 2009. The Listeria transcriptional landscape from saprophytism to virulence. Nature 459:950–6.

41. Mattick JS. 2002. Type IV pili and twitching motility. Annu Rev Microbiol 56:289–314.

42. Dwyer DJ, Belenky PA, Yang JH, MacDonald IC, Martell JD, Takahashi N, Chan CT, Lobritz MA, Braff D, Schwarz EG, Ye JD, Pati M, Vercruysse M, Ralifo PS, Allison KR, Khalil AS, Ting AY, Walker GC, Collins JJ. 2014. Antibiotics induce redox-related physiological alterations as part of their lethality. Proc Natl Acad Sci U S A 111:E2100–9.

43. Lobritz MA, Belenky P, Porter CB, Gutierrez A, Yang JH, Schwarz EG, Dwyer DJ, Khalil AS, Collins JJ. 2015. Antibiotic efficacy is linked to bacterial cellular respiration. Proc Natl Acad Sci U S A 112:8173–80.

44. Kohanski MA, Dwyer DJ, Hayete B, Lawrence CA, Collins JJ. 2007. A common mechanism of cellular death induced by bactericidal antibiotics. Cell 130:797–810.

45. Belenky P, Ye Jonathan D, Porter Caroline BM, Cohen Nadia R, Lobritz Michael A, Ferrante T, Jain S, Korry Benjamin J, Schwarz Eric G, Walker Graham C, Collins James J. 2015. Bactericidal Antibiotics Induce Toxic Metabolic Perturbations that Lead to Cellular Damage. Cell Reports 13:968–980.

46. Dingsdag SA, Hunter N. 2018. Metronidazole: an update on metabolism, structure-cytotoxicity and resistance mechanisms. J Antimicrob Chemother 73:265–279.

47. Knippel RJ, Wexler AG, Miller JM, Beavers WN, Weiss A, de Crecy-Lagard V, Edmonds KA, Giedroc DP, Skaar EP. 2020. Clostridioides difficile Senses and Hijacks Host Heme for Incorporation into an Oxidative Stress Defense System. Cell Host Microbe doi: 10.1016/j.chom.2020.05.015.

48. Sambrook J, Russell DW. 2001. Molecular cloning : a laboratory manual. Cold Spring Harbor Laboratory, Cold Spring Harbor, N.Y.

49. Purdy D, O’Keeffe TA, Elmore M, Herbert M, McLeod A, Bokori-Brown M, Ostrowski A, Minton NP. 2002. Conjugative transfer of clostridial shuttle vectors from Escherichia coli to Clostridium difficile through circumvention of the restriction barrier. Mol Microbiol 46:439–52.

50. Hofmann JD, Otto A, Berges M, Biedendieck R, Michel A-M, Becher D, Jahn D, Neumann-Schaal M. 2018. Metabolic Reprogramming of Clostridioides difficile During the Stationary Phase With the Induction of Toxin Production. Frontiers in Microbiology 9.

51. Cartman ST, Kelly ML, Heeg D, Heap JT, Minton NP. 2012. Precise manipulation of the Clostridium difficile chromosome reveals a lack of association between the tcdC genotype and toxin production. Appl Environ Microbiol 78:4683–90.

52. Sebaihia M, Wren BW, Mullany P, Fairweather NF, Minton N, Stabler R, Thomson NR, Roberts AP, Cerdeno-Tarraga AM, Wang H, Holden MT, Wright A, Churcher C, Quail MA, Baker S, Bason N, Brooks K, Chillingworth T, Cronin A, Davis P, Dowd L, Fraser A, Feltwell T, Hance Z, Holroyd S, Jagels K, Moule S, Mungall K, Price C, Rabbinowitsch E, Sharp S, Simmonds M, Stevens K, Unwin L, Whithead S, Dupuy B, Dougan G, Barrell B, Parkhill J. 2006. The multidrug-resistant human pathogen Clostridium difficile has a highly mobile, mosaic genome. Nat Genet 38:779–86.

53. Dupuy B, Sonenshein AL. 1998. Regulated transcription of Clostridium difficile toxin genes. Mol Microbiol 27:107–20.

54. van Eijk E, Anvar SY, Browne HP, Leung WY, Frank J, Schmitz AM, Roberts AP, Smits WK. 2015. Complete genome sequence of the Clostridium difficile laboratory strain 630Deltaerm reveals differences from strain 630, including translocation of the mobile element CTn5. BMC Genomics 16:31.

55. Dannheim H, Riedel T, Neumann-Schaal M, Bunk B, Schober I, Sproer C, Chibani CM, Gronow S, Liesegang H, Overmann J, Schomburg D. 2017. Manual curation and reannotation of the genomes of Clostridium difficile 630Deltaerm and C. difficile 630. J Med Microbiol 66:286–293.

